# Social hierarchy regulates ocular dominance plasticity in adult male mice

**DOI:** 10.1101/579169

**Authors:** Jenny Balog, Franziska Hintz, Marcel Isstas, Manuel Teichert, Christine Winter, Konrad Lehmann

## Abstract

We here show that social rank, as assessed by competition for a running wheel, influences ocular dominance plasticity in adult male mice. Dominant animals showed a clear ocular dominance shift after four days of MD, whereas their submissive cage mates did not. NMDA receptor activation, reduced GABA inhibition, and serotonin transmission were necessary for this plasticity, but not sufficient to explain the difference between dominant and submissive animals. In contrast, prefrontal dopamine concentration was higher in dominant than submissive mice, and systemic manipulation of dopamine transmission bidirectionally changed ocular dominance plasticity. Thus, we could show that a social hierarchical relationship influences ocular dominance plasticity in the visual cortex via higher-order cortices, most likely the medial prefrontal cortex. Further studies will be needed to elucidate the precise mechanisms by which this regulation takes place.

## 1. Introduction

Dominance and submissiveness occur in all areas of life, not only among people in school, work and other social situations, but in all group living animals (Stears et al. 2014). The physiological background of this social condition is being studied in many vertebrate (Morgan et al. 2002; Desjardins and Fernald 2008; Kar et al. 2017, Jetz and Rubenstein 2011), and invertebrate species (Sbragaglia et al. 2017). A social hierarchy entails a dominant-submissive relationship between all individual pairs of animals within the group. As a result, most animals will experience defeat and subordination frequently. In rodents, this experience has been shown to induce stress (Blanchard et al. 1995), compromise mental health (Prabhu et al. 2018), and impair learning (Goeckner et al. 1973; Spritzer et al. 2004). More recent studies in pairs of mice have confirmed that, indeed, learning ability deteriorates in submissive animals, which is not, however, directly due to stress (Fitchett et al. 2005; Colas-Zelin et al. 2012; Matzel et al. 2017).

Whereas an impact of social dominance or submissiveness on behavioural learning has thus been well established, an effect on more basal cortical plasticity has not yet been investigated. We have recently shown that paired, in contrast to individual, housing of adult mice reinstated ocular dominance plasticity (ODP, Balog et al. 2014), i.e. the propensity of the primary visual cortex (V1) to shift its responsiveness towards the open eye when one eye is experimentally closed. Whereas in female mice, which are not aggressive and do not establish a clear hierarchy, this effect was seen irrespective of the available space, it was only present in both male mice of a pair if they disposed of a large arena. In a standard cage, only one of the two would show plasticity.

The assumption was obvious that the difference in plasticity was due to social dominance, the cramped space forcing the males to arrange their relationship differently than in the large arena. In this study, we tested this hypothesis and went on to elucidate the mechanisms by which social status regulates ocular dominance plasticity in male mice. In addition to biochemical factors acting within the visual cortex, we found dopamine, acting most probably in the prefrontal cortex, to play a decisive role.

## 2. Material and Methods

### 2.1 Animals and housing conditions

Fully adult (> postnatal day 133) male C57BL/6J mice were housed in groups of 2–3 in transparent standard Makrolon cages (16.5 × 22.5 cm^2^, minimally enriched with cotton rolls and nest material) on a 12 hr light/ dark cycle, with food and water available ad libitum. The experiment started with joining two animals out of different groups into one standard cage. Beginning immediately, the animals got access to a running wheel for 1.5h per day during three days before and after MD, to assess dominance behaviour (see below). In the arena condition, two mice together were kept six hours per day in a square, featureless arena with a side length of 72 cm, and were then put back into their single cages. All groups, including the control group animals, were treated identically as described above. In our institution, animal housing is constantly supervised by veterinaries from the state of Thuringia, Germany. All experimental procedures have been performed according to the German Law on the Protection of Animals and the corresponding European Communities Council Directive of 2010 (2010/63/EU), and were approved by the Thüringer Landesamt für Lebensmittelsicherheit und Verbraucherschutz (Thuringia State Office for Food Safety and Consumer Protection) under the registration number 02-036/15.

### 2.2 Behavioural analysis

Male mice are territorial and will form a dominance hierarchy when forced to live together (Kappel et al. 2017). For obvious animal welfare reasons, we could not maintain pairings in which escalating fighting was observed. In peaceful mice, however, dominance hierarchy is hard to determine after its initial agonistic establishment. We therefore developed several behavioural criteria for the reliable determination of social dominance.

First, whenever possible, we arranged pairings in which one of the mice lacked the vibrissae, since they were likely the victims of so-called “barbering” by dominant mice in their group of origin (Sarna et al. 2000), and therefore possible predisposed for becoming submissive again.

Second, we routinely assessed competition for a limited resource in all pairings. For mice, use of a running wheel is so desirable that they will even access it in the wild (Meijer and Robbers 2014). Indeed, when the pairing cage was furnished with just one running wheel, it became quickly apparent which mouse asserted privileged access to it. Although it has been claimed that running wheel use during MD can by itself elicit OD plasticity (Kalogeraki et al. 2014), we have previously conclusively shown that locomotion or running wheel use has no such effect during short-term MD of single or paired housed mice (Balog et al. 2014). Potential reasons for the discrepancy will be dealt with in the discussion.

Thus, we provided a running wheel to both animals together for three days before and after monocular deprivation for 1h per day. Turns run by each mouse were counted automatically, converted into m/h and averaged across days. In order to ensure that differential use of the running wheel was not due to differing interest, each mouse was subsequently put alone for 1/2 h into a running wheel cage, and turns were counted. Running wheel use alone and in pairs could then be quantitatively compared.

Third, in order to corroborate that running wheel use is a reliable measure of social dominance, we also performed a behavioural analysis by counting behaviours defined in an ethogram based on a published template (Olsson and Sherwin 2006). The following behaviours were counted while the mice were together in the running wheel cage:

Attack – jumping at or chasing the other mouse, biting, kicking, wrestling
Flight – avoidance of contact, direct withdrawal from the other mouse
Head sniff – sniffing directed to the head (mostly nose) of the other mouse
Anal sniff – sniffing directed to the anus of the other mouse
Social grooming – licking and nibbling the other mouse at different areas of the body
Allogrooming – more intense grooming: Vigorous licking with a higher incidence of teeth hair pulling. The other mouse is usually motionless; otherwise it comes to a fight

### 2.3 Monocular Deprivation

MD was always performed after the first imaging session according to published protocols (Gordon and Stryker 1996; Lehmann et al. 2012). Briefly, mice were anaesthetized by 2% isoflurane in a mixture of 1:1 N2O and O2. Before the imaging session, animals received one injection of carprofen (4 mg/kg, s.c.). Lid margins of the right eye were trimmed and an antibiotic ointment was applied before the eye was sutured shut. All animals were checked daily to ensure that the sutured eye remained closed during the 4 days of MD. Over the following days animals received a daily administration of carprofen (0.12 mg/mouse, s.c.).

### 2.4 Drug administration

#### WAY-100635

To investigate if serotonin plays a role for the restored OD plasticity, we administrated 1 mg/kg i.p. of the 5-HT_1A_-receptor antagonist WAY-100635, diluted in saline (Fletcher et al. 1996; Forster et al. 1995).

#### Diazepam

To test if changes in the activity of GABA are necessary to trigger adult OD plasticity, we treated the animals with a daily i.p. injection of 1 mg/kg diazepam (Stodieck et al. 2014).

#### CPP

To find out the role of the N-methyl-D-aspartate (NMDA)-receptor on ocular dominance plasticity, we administrated the NMDA receptor-Antagonist (R,S)-3-(2-carbooxypiperazin-4-yl)propyl-1-phosphonic (CPP) at a daily dose of 12–15 mg/kg i.p. in saline (Sato and Stryker 2008; Villarreal et al. 2002).

#### Zuclopenthixol

To investigate a possible role of cortical dopamine transmission in OD plasticity, we treated the animals with the dopamine receptor (D1 and D2) antagonist Zuclopenthixol at a low dose that reduces aggression without motor effects (Manzaneque and Navarro 1999). It was diluted in methylcellulose (15 % in aqua bidest) and 15 % ethanol and injected i.p. every 24 hours at a dose of 0.2 mg/kg.

#### Methylphenidate

To enhance cortical dopamine transmission, we administrated the dopamine reuptake inhibitor methylphenidate-hydrochloride at a low dose that increases dopamine transmission only in the mPFC but not the striatum (Koda et al. 2010). It was diluted in 0.9% NaCl and injected four times a day at a dose of 3 mg/kg.

### 2.5 Optical Imaging of intrinsic signals

#### 2.5.1 Mouse preparation for optical imaging

Animals were initially anesthetized with 4% isoflurane in a mixture of 1 : 1 O_2_/N_2_O and placed on a heating blanket (37.5 °C) for maintaining body temperature. They then received injections of chlorprothixene (0.2 mg/mouse, i.m.), atropine (0.3 mg/mouse, s.c.), dexamethasone (0.2 mg/mouse, s.c.) and carprofen (0.12 mg/mouse, s.c.). Inhalation anesthesia was applied through a plastic mask and maintained at 1.5% during the surgical intervention and 0,5% during data recording. The animal was fixed in a stereotaxic frame using non-crush ear bars. The skin above the left hemisphere was removed to expose the visual cortex. This exposed area was covered with 2.5% agarose in saline and sealed with a glass coverslip. Cortical responses were always recorded through the intact skull. Immediately after the first imaging session the skin was resutured and MD was performed. After that the animals were returned to their standard cage which was placed on a heating plate overnight. Mice were checked every 15 minutes until the righting reflex was positive. Before the next imaging session, the skin and the eye were reopened and imaging was performed as described above.

#### 2.5.2 Imaging of visual cortex

The imaging method of temporally encoded maps was originally described by Kalatsky and Stryker (2003). Using a Dalsa 1M30 CCD camera (Dalsa, Waterloo, Canada) with a 135 mm x 50 mm tandem lens configuration (Nikon, Inc. Melville, NY), we imaged an area of 4.6 x 4.6 mm^2^. The surface vascular pattern was visualized by green illumination (550 ± 2 nm). Thereafter, focusing 600 μm below the pial surface, intrinsic signals were obtained via illumination with red light (610 ± 2 nm). Frames were acquired at a rate of 30 Hz and temporally averaged to 7.5 Hz. The 1024×1024 pixel images were spatially binned to a 512 x 512 resolution. Horizontal drifting bars (2° wide) were presented at a temporal frequency of 0.125Hz and were adjusted so that they only appeared in the binocular visual field of the recorded left hemisphere (−5° to +15° azimuth). Thus, the stimulus was repeated for about 35 times during one presentation. Cortical responses were extracted by Fourier analysis at the stimulation frequency and converted into amplitude and phase maps using custom software (Kalatsky and Stryker 2003). Ocular dominance indices (ODIs) were calculated from activity maps as described previously (Cang et al. 2005; Lehmann and Löwel 2008). Briefly, within a region of interest, all pixels above a threshold at 30 % of peak amplitude were used, and OD was calculated for each pixel as (contra − ipsi) / (contra + ipsi), and averaged across all selected pixels.

### 2.6 Post-mortem HPLC

After the second session of optical imaging, the scalp was sutured and the animals were allowed to re-awake. The following day, they were again transferred to their respective condition for another three days. After that, the animals were killed by cervical dislocation, the brains were quickly dissected and frozen immediately at −40 °C. Neurotransmitter contents were measured using high performance liquid chromatography (HPLC). Micropunches were taken from 1 mm mPFC slices at 2.34 from Bregma and 1 mm V1 slices at −3.28 from Bregma (Paxinos and Franklin 2012). These brain samples were homogenized by ultrasonication in 20 vol of 0.1 N perchloric acid at 4 °C immediately after collection. A total of 100 μl of the homogenate was added to equal volumes of 1 N sodium hydroxide for measurement of protein content. The remaining homogenate was centrifuged at 17000 g and 4 °C for 10min. Supernatants were used for immediate measurement. The levels of monoamines (DA and 5-HT) and their metabolites (DOPAC, and 5-HIAA) were measured by HPLC with electrochemical detection as previously described (Felice et al. 1978; Sperk et al. 1981; Sperk 1982; Enard et al. 2009; Winter et al. 2009; Giovanoli et al. 2013). Briefly, the perchloric acid extracts were separated on a column (Prontosil 120-3-C18-SH; length 150 mm, inner diameter 3 mm; Bischoff Analysentechnik und- geräte GmbH, Leonberg, Germany) at a flow rate of 0.55 ml/min. The mobile phase consisted of 80 mM sodium dihydrogene phosphate, 0.85 mM octane-1-sulfonic acid sodium salt, 0.5 mM ethylenediaminetetraacetic acid disodium salt, 0.92 mM phosphoric acid and 4% 2-propanol (all chemicals Merck KGaA, Darmstadt, Germany). Monoamines were detected using an electrochemical detector (41 000, Chromsystems Instruments & Chemicals GmbH, Munich, Germany) at an electrode potential of 0.8 V. For calibration, 0.1 M perchloric acid containing 0.1 mM DOPAC, 5-HIAA and 5-HT and 1 mM DA was injected into the HPLC system before and after sample analysis. Sample analysis was performed based on peak areas using a computer-based chromatography data system (CSW 1.7, DataApex Ltd, Prague, Czech Republic) in relation to the mean of the applied calibration solutions.

### 2.7 Retrograde fluorescent tracing

Mice were prepared for optical imaging (see above), and one map of absolute retinotopy was acquired in the left hemisphere. The map was overlaid over an image of the blood vessel pattern. Trepanations were performed over the parts of the visual cortex that represented the upper and the lower visual field. A green retrograde tracer (CTb-488) was injected into the rostral, a red tracer (Ctb-594) into the caudal visual cortex. Then, the scalp was sutured, and the animal was allowed to wake up.

One week later, the animal was perfused with 0.1M PBS, followed by 4% PFA in PBS. The brain was dissected and frozen after cryoprotection. 50μm slices were taken on a vibratome, counterstained with DAPI and coverslipped with Mowiol.

### 2.8 Statistical analysis

All group comparisons were initially subjected to two-way ANOVA, with repeated measures where appropriate, in order to check for main effects. Optical imaging data were further analyzed by post hoc two-tailed student’s t-tests, with paired t-tests for before–after comparisons and unpaired t-tests for between-group comparisons. The resulting p-values were Bonferroni corrected. The levels of significance were set as *: p ≤ 0.05; **: p ≤ 0.01; ***: p ≤ 0.001. Data are presented as means and standard error of the mean (s.e.m.).

## 3. Results

### 3.1 Running wheel use reflects and induces social hierarchy in mice

Having previously shown that of a pair of male mice housed in a standard cage, only one will show ocular dominance (OD) plasticity after 4d of MD (Balog et al. 2014), we here wished to test our hypothesis that this difference is due to a social hierarchy. To this end, we established a novel dominance test: the mice got access to a running wheel, both together (1h/day) and alone (1/2h/day), three days before and three days during MD. Figure 1A shows running wheel use of mice classified as “dominant” (grey columns) or “submissive” (white columns) both while together and while alone. In each pair, one mouse each was assigned to either group based on these measures. While being together in the running wheel cage, dominant mice accessed the running wheel significantly more (1235.83 ± 106.63 m/h) than their submissive cagemates (458.78 ± 63.94 m/h, p ≤ 0.001, t-test; n = 12). When the animals were alone, this difference did not entirely disappear (p ≤ 0.05, t-test), but was strongly reduced, which was due to submissive (923.36 ± 132.69 m/h, p ≤ 0.001 compared to paired) – not dominant (1331.96 ± 148.20 m/h, n.s. compared to paired) – mice drastically increasing their use of the wheel. This shows that mice classified as submissive did not lack interest in the wheel, but refrained from its use because the dominant mouse was present.

**Figure 1:**
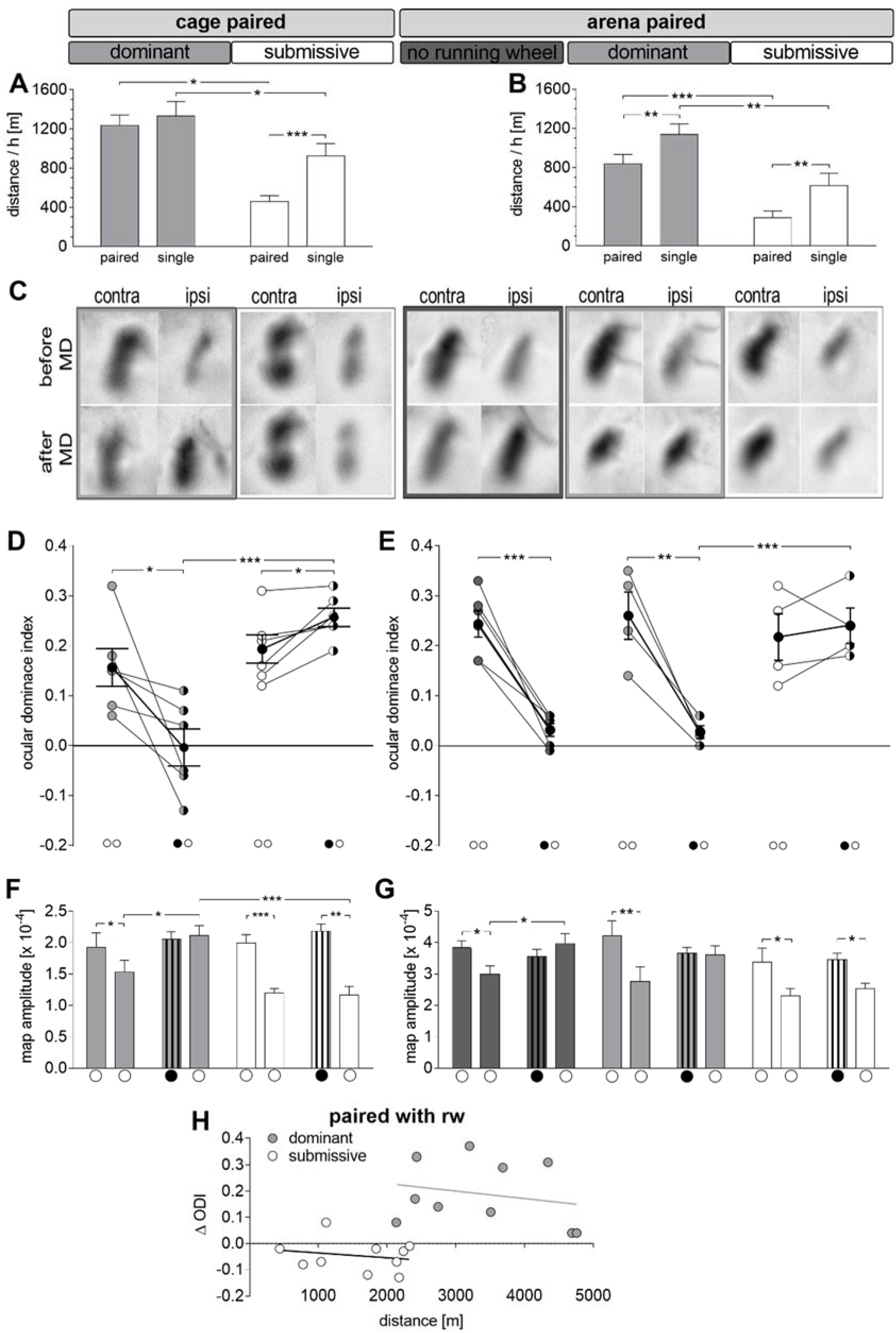
Social dominance determines ocular dominance plasticity. Both in a cage (A) and in an arena (B), a running wheel is predominantly used by one of two male animals. When alone, the less active animals significantly increase running. (C) Amplitude maps obtained by optical imaging of intrinsic signals are shown. Whereas stronger activities are always elicited by stimulation of the contralateral than the ipsilateral eye before MD, this difference is lost or even reversed in dominant mice or arena mice without a running wheel. (D) In a cage, dominant mice showed OD plasticity after 4d of MD, whereas submissive mice did not. (E) When housed in the arena, all mice showed full plasticity, but once social hierarchy was induced by the presence of a running wheel, OD plasticity disappeared in submissive animals. Each symbol represents the ODI of an individual animal, horizontal lines show the group mean. Full symbols represent control measurements, half symbols measurement after MD. White circles show open eyes, black circles closed contralateral eyes. Dominant animals are shown as grey, submissive animals as white symbols. (F, G) V1 activity elicited by contra- or ipsilateral eye stimulation shows that OD plasticity was achieved by open-eye potentiation both in cage dominant mice and in arena mice without running wheel, whereas there was no significant result in dominant mice in the arena with running wheel. (H) Running activity had no influence on OD plasticity (ODI before MD – ODI after MD) neither in submissive nor in dominant mice.

Ethological data corroborated this interpretation. In all cases, mice classified as dominant because of the running wheel data performed attacks at a much higher rate – usually exclusively – than the submissive mice, which correspondingly showed flight behaviour. Frequently, submissive mice were chased from the wheel by their dominant cagemates. In all cases, these observations confirmed the ranking derived from wheel use.

We have previously shown that paired male mice in a featureless arena differ significantly in their behaviour from pairs in a cage, and both show OD plasticity at an indistinguishable level (Balog et al. 2014). This indicates that social hierarchy is not established in the absence of a limited resource. In order to test this assumption, we introduced a running wheel into an otherwise bare arena. Indeed, its use by the two mice was disbalanced, with one animal occupying it most of the time and defending the access violently, yielding very similar results to the cage condition (dominant paired 838.32 ± 96.27 m/h vs. submissive paired 288.01 ± 68.58 m/h; p ≤ 0.001, t-test; dominant single 1137.93 ± 108.55 m/h vs. submissive single 614.87 ± 126.99 m/h; p ≤ 0.01, t-test; n = 8). The mice were accordingly classified as “dominant” and “submissive” (Fig. 1B). We proceeded to test whether the behavioural differences would result in different brain plasticity.

### 3.2 The dominance status of adult male mice regulates ocular dominance plasticity

Ocular dominance was measured in the same cage mice before and after 4d MD via optical imaging (Fig. 1C, D). At control imaging, darker activity spots were always obtained by stimulation of the contralateral than ipsilateral eye (Fig. 1C), and dominant (0.16 ± 0.04) and submissive (0.19 ± 0.03) mice had on average similar OD values (p = 0.45, t-test). After 4d of MD, however, there was a significant OD shift towards the open ipsilateral eye in dominant mice (−0.003 ± 0.04, n = 6), whereas the ODI even increased in submissive animals (0.26 ± 0.02, n = 6). ANOVA with repeated measures revealed significant influences of social dominance (F_1.10_ = 22.542, p ≤ 0.001) and its interaction with MD (F_1.10_ = 12.483, p ≤ 0.01). This allowed us to do post-hoc paired t-tests, which confirmed significant (p ≤ 0.05) shifts due to MD in both the dominant and submissive groups, with the effect that ODIs were highly significantly different between dominant and submissive mice after MD (p ≤ 0.001).

To our surprise, the same difference in plasticity was seen in arena mice after the introduction of a running wheel (Fig. 1C, E). First, we confirmed our previous finding that in a bare arena, both male animals show full OD plasticity. Indeed, the ODI shifted towards lower values in all animals, on average from 0.24 ± 0.03 to 0.03 ± 0.01 after MD (p ≤ 0.001, paired t-test; n = 6). If in the same situation a running wheel was present, leading to the establishment of a social hierarchy (see above), OD plasticity likewise differed between the two animals of a pair, indicated by significant effects of MD (F_1.6_=12.798, p<0.05) and its interaction with social dominance (F_1.6_=18.871, p ≤ 0.01). In dominant mice, there was a significant OD shift (0.26 ± 0.05 to 0.03 ± 0.01, n = 4; p ≤ 0.01, paired t-test), which was absent in submissive mice (0.22 ± 0.05 to 0.24 ± 0.04, n = 4; p = 0.58, paired t-test). Accordingly, these animals had significantly higher OD values after MD than dominant animals (p ≤ 0.001, t-test). Thus, the introduction of a running wheel for only one week altogether induced a social hierarchy that resulted in differential ocular dominance in male mice. Since behavioural and physiological data were similar in cage and arena mice with a running wheel, the respective dominant and submissive groups were pooled for the later biochemical investigations.

Running by itself has been indicated to induce OD plasticity in one study (Kalogeraki et al. 2014). As we have employed running wheel use – albeit only for a short time per day – to assess social dominance, one might argue that the difference in OD plasticity between dominant and submissive mice could trivially be explained by their different amount of running. Although we have already refuted this assumption in our previous study (Balog et al. 2014), we wished to further check this possibility by correlating the total running distance of each animal with its plasticity, i.e. the difference between ODIs before and after MD (Fig. 1H). Cage and open field mice were pooled. Although, as expected, all submissive mice are in the lower left quadrant, and all dominant ones in the upper right, which would obviously result in a positive and significant correlation, no such correlation is observed within each group (dominant: r=-0.22, p=0.54; submissive: r=-0.21, p=0.57, regression analysis). Note, too, the overlap at approx. 2200m running distance, with does not entail an overlap in plasticity. Thus, brief running for 1.5h per day during four days of MD does not induce OD plasticity in adult male mice.

We wondered whether the observed OD shifts in dominant animals were due to an increase in open-eye activity (so-called adult plasticity) or a decrease in closed-eye activity (juvenile plasticity). In caged pairs, the OD shift was achieved by strengthening of the ipsilateral (open)-eye input (Fig. 1F, ipsi: [1.53 ± 0.19] x 10^−4^ to [2.11 ± 0.16] x 10^−4^; p ≤ 0.05, t-test; contra: [1.93 ± 0.23] x 10^−4^ to [2.06 ± 0.12] x 10^−4^; p = 0.5; t-test). The response amplitudes of contralateral (closed) and ipsilateral (open) eyes of the submissive mice did not change after 4d MD (ipsi: [1.2 ± 0.07] x 10^−4^ to [1.17 ± 0.13] x 10^−4^; p = 0.8, t-test; contra: [1.99 ± 0.14] x 10^−4^ to [2.18 ± 0.11] x 10^−4^; p = 0.2; t-test).

In the arena (Fig. 1G), OD plasticity was also of the adult type in the control condition, with a significant increase of the open ipsilateral eye response after 4d of MD ([2.98 ± 0.72] x 10^−4^ to [3.97 ± 0.94] x 10^−4^; p ≤ 0.05, t-test) and no change of the contralateral eye response ([3.83 ± 0.26] x 10^−4^ to [3.55 ± 0.29] x 10^−4^; p = 0.5; t-test). In the presence of a running wheel, changes in both open and closed eye amplitudes were non-significant in dominant animals (ipsi: [2.76 ± 0.46] x 10^−4^ to [3.61 ± 0.28] x 10^−4^; p = 0.3, t-test; contra [4.18 ± 0.49] x 10^−4^ to [3.66 ± 0.18] x 10^−4^; p = 0.5; t-test). The response amplitudes of contralateral and ipsilateral eyes of the submissive mice did not differ after 4d MD (ipsi: [2.36 ± 0.24] x 10^−4^ to [2.53 ± 0.17] x 10^−4^; p = 0.6, t-test; contra [3.38 ± 0.44] x 10^−4^ to [3.45 ± 0.2] x 10^−4^; p = 0.9; t-test).

### 3.3 Ocular dominance plasticity in adult dominant male mice is dependent on the activation of the serotonin receptor 5-HT_1A_

It has already been shown that adult ocular dominance plasticity can be restored by an increased serotonin transmission (Maya Vetencourt et al. 2008, 2011), and that the effects of enriched environment and social experience, which also induce OD plasticity, are mediated by this transmitter (Baroncelli et al. 2010; Balog et al. 2014). We therefore wanted to check if serotonin plays a role for the reinstated plasticity of socially dominant mice. After the second optical imaging session the mice were returned to their respective housing conditions for another three days, whereupon they were sacrificed for post-mortem HPLC. Figure 2A shows that the contents of serotonin (5-HT) and its metabolite 5-hydroxy-indole-acetic acid (5-HIAA) were not different between dominant and submissive paired mice (dominant 5-HT: 8.38 nmol/mg protein ± 1.38 vs. submissive 5-HT: 8.41 nmol/mg protein ± 1.83; p = 1.0; t-test; dominant 5-HIAA: 2.93 nmol/mg protein ± 0.65 vs. submissive 5-HIAA: 2.92 nmol/mg protein ± 0.84; p = 1.0; t-test; n = 9). The 5-HT turnovers (5-HIAA/5-HT-ratio), too, were indistinguishable (dominant: 0.35 ± 0.03, submissive: 0.36 ± 0.05, p=0.91, t-test). Data of cage mice and mice from an arena with running wheel were pooled for this analysis (see above). This result was surprising, as it effectively excludes serotonin as an inductor of OD plasticity in our present model.

**Figure 2:**
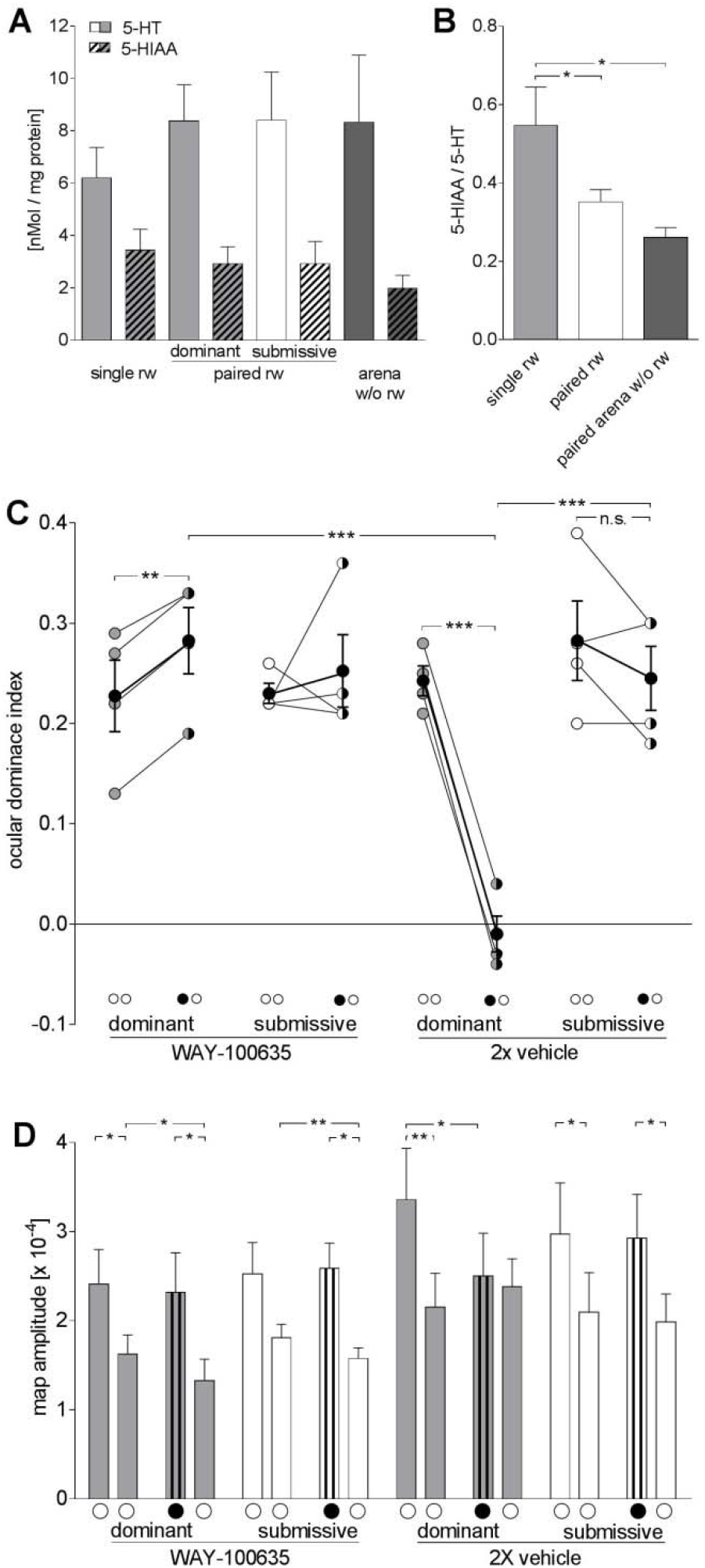
Adult ocular dominance plasticity induced by social experience requires 5-HT_1A_ receptor activation. (A) The contents of serotonin (5-HT) and its metabolite 5-HIAA were determined by HPLC in the visual cortices. There was no difference in 5-HT or 5-HIAA content between paired dominant and submissive cage mice. (B) 5-HT turnover (5HIAA / 5-HT ratio) was significantly higher in single cage mice than in both paired mice groups. (C) Ocular dominance indices show that there is no ocular dominance plasticity in dominant paired cage mice treated with the 5HT_1A_ receptor antagonist WAY-100635, whereas full plasticity was observed in vehicle-injected animals. (D) Response amplitudes show an unexpected decrease in ipsilateral eye responses in WAY-100635-treated animals, and an equally unexpected decrease in contralateral eye responses in dominant vehicle animals. All conventions are as in Fig. 1.

We therefore wondered whether it might at least be a permissive factor. To answer this question, we compared the contents of 5-HT and 5-HIAA in pooled dominant and submissive mice to both single-housed mice (which show no plasticity) and arena mice (which show serotonin-dependent plasticity, Balog et al. 2014). 5-HT and 5-HIAA contents (Fig. 2A) were highly variable in our hands and precluded any conclusion: Both single mice (5-HT: 6.2 ± 1.17 nmol/mg protein and 5-HIAA: 3.45 ± 0.79nmol/mg protein; n = 5), and paired mice in an arena without a running wheel (5-HT: 8.32 ± 2.56nmol/mg protein and 5-HIAA: 1.99 ± 0.49 nmol/mg protein; n = 4) showed similar values as caged pairs. However, the 5-HT turnover (5-HIAA/5-HT-ratio) was significantly affected by housing condition (F_2,24_ = 5.09, p < 0.05, ANOVA), as it was lower in both pair-housed groups than in single-housed animals (single cage: 0.55 ± 0.1, n = 5, paired cage with rw: 0.35 ± 0.03, n = 18, p < 0.05, paired arena w/o rw: 0.26 ± 0.03, n = 4, p < 0.05, Tukey test). This finding suggests that 5-HT transmission is not sufficient to induce OD plasticity in dominant male mice, but is necessary as a permissive factor.

To test this interpretation, we treated animals with WAY-100635, the antagonist of the 5-HT_1A_-receptor (Fig. 2C). Indeed, this intervention inhibited OD plasticity in dominant paired cage animals (0.23 ± 0.04 to 0.28 ± 0.03, n = 4), resulting even in a significantly higher ODI after 4d of MD. ANOVA showed significant effects of social dominance (F_1.12_ = 6.35, p ≤ 0.05), MD (F_1.12_ = 17.752, p ≤ 0.001) and treatment (F_1.12_ = 4.797, p ≤ 0.005), as well as all of their interactions (all p ≤ 0.01). The ODI did not change in the submissive paired cage animals during MD (0.23 ± 0.01 to 0.25 ± 0.04, n = 4, p = 0.6, paired t-test). In the saline control group, the ODI of the dominant paired cage control animals was significantly decreased after 4d MD (0.24 ± 0.02 to −0.01 ± 0.02, n = 4; p ≤ 0.001, paired t-test). As expected, the ODI of the submissive paired cage mice did not change (0.28 ± 0.04 to 0.25 ± 0.03, n = 4; p = 0.3, paired t-test). These results confirm that 5-HT_1A_ receptor activation is required to mediate the enhanced OD plasticity in adult dominant paired cage mice.

Analysis of the response amplitudes (Fig. 2D) confirmed that cortical activation by the contralateral eye remained significantly stronger than by the ipsilateral eye in all conditions where OD plasticity was absent. Apart from the factor “eye” and its interactions with the other variables, no other factor had any effect on the amplitudes, according to ANOVA. Somewhat surprisingly, ipsilateral eye responses decreased significantly in both dominant and submissive animals (dominant: contra: [2.41 ± 0.39] x 10^−4^ to [2.32 ± 0.44] x 10^−4^, p = 0.7, ipsi: [1.63 ± 0.21] x 10^−4^ to [1.33 ± 0.24] x 10^−4^, p ≤ 0.05; submissive: contra: [2.53 ± 0.35] x 10^−4^ to [2.59 ± 0.28] x 10^−4^, p = 0.9; ipsi: [1.81 ± 0.15] x 10^−4^ to [1.58 ± 0.12] x 10^−4^, p ≤ 0.01; all comparisons by t-test).

Also to our surprise, there was a significant decline in contralateral deprived-eye responses to open-eye levels in dominant vehicle animals (contra: [3.36 ± 0.57] x 10^−4^ to [2.5 ± 0.48] x 10^−4^, p ≤ 0.05; ipsi: [2.15 ± 0.38] x 10^−4^ to [2.39 ± 0.31] x 10^−4^, p = 0.15 t-test). This juvenile plasticity is at conspicuous variance to the adult plasticity observed in completely untreated animals.

### 3.4 Ocular dominance plasticity in socially dominant male mice is regulated by cortical inhibition and long term potentiation

Restored OD plasticity in fully adult mice is frequently associated with a decrease in cortical GABA release, and can be blocked by diazepam administration (Hanover et al. 1999; Harauzov et al. 2010; Huang et al. 1999; Maya Vetencourt et al. 2011; Sale et al. 2007). It has repeatedly been shown, too, that the initial phase of both juvenile and adult OD plasticity consists of NMDA-receptor-dependent Hebbian LTP and LTD (Sawtell et al. 2003; Sato and Stryker 2008; Ranson et al. 2012). To check whether the same plasticity mechanisms are at work in socially dominant male mice, we treated experimental groups of mice with either diazepam or the NMDA receptor blocker CPP.

Figure 3A shows that OD plasticity in dominant paired cage mice was abolished by both interventions, diazepam (0.25 ± 0.04 to 0.24 ± 0.02, n = 4, p = 0.7, t-test) and CPP (0.27 ± 0.03 to 0.23 ± 0.05, n = 5, p = 0.3, t-test). As usual, the submissive paired cage animals did not show any OD plasticity, neither under diazepam treatment (0.24 ± 0.02 vs. 0.23 ± 0.04, n = 4, p = 0.7, t-test), nor under CPP treatment (0.29 ± 0.02 vs. 0.27 ± 0.02, n = 5, p = 0.08, t-test). As the vehicle group was included into the ANOVA, there were significant effects of MD (diazepam: F_1.12_ = 11.084, p ≤ 0.001, CPP: F_1.14_ = 31.116, p ≤ 0.001) and treatment (diazepam: F_1.12_ = 11.64, p ≤ 0.01; CPP: F_1.14_ = 15.628, p ≤ 0.001) for both interventions, together with all interventions. By post-hoc testing, these effects could be attributed to the fact that in the vehicle group, a strong OD shift was found in dominant paired cage animals (0.22 ± 0.03 to 0.01 ± 0.03, n = 4, p ≤ 0.001, t-test), which made the values significantly different from those in the corresponding treatment groups (p ≤ 0.001 and p ≤ 0.01, respectively). This shift was here again due to an increase in open-eye responses (Fig. 3B, [2.74 ± 0.18] x 10^−4^ vs. [3.06 ± 0.46] x 10^−4^; p ≤ 0.05; t-test), whereas the response of the contralateral eyes stayed the same. The animals receiving vehicle once per day thus resemble untreated animals. The ODI of submissive control animals did not change after 4d of MD.

**Figure 3:**
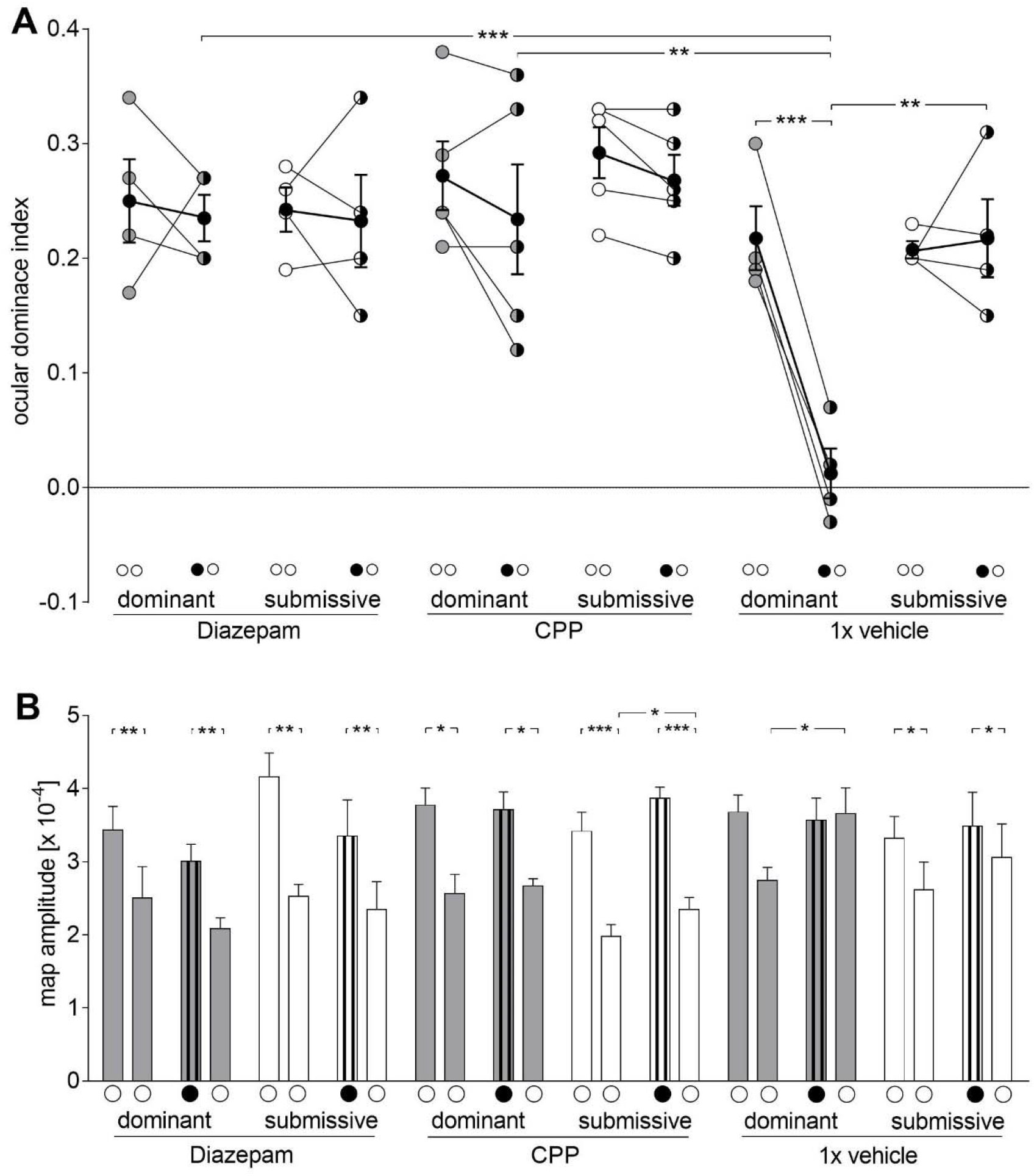
Cortical inhibition and long term potentiation are involved in social dominance-induced OD plasticity. (A) Both diazepam and CPP blocked OD plasticity in socially dominant male mice. (B) Contralateral eye responses remained higher than ipsilateral eye responses in all treated groups, but there was a significant increase of ipsilateral eye responses in dominant vehicle mice. All conventions are as in Fig. 1.

In all diazepam-treated groups, V1 activity elicited by contra- or ipsilateral eye stimulation remained unchanged in dominant (contra: [3.44 ± 0.32] x 10^−4^ vs. [3.01 ± 0.23] x 10^−4^; p = 0.3; ipsi: [2.5 ± 0.43] x 10^−4^ vs. [2.08 ± 0.15] x 10^−4^; p = 0.31; t-test) and submissive animals (contra: [4.16 ± 0.32] x 10^−4^ vs. [3.35 ± 0.49] x 10^−4^; ipsi: [2.52 ± 0.17] x 10^−4^ vs. [2.34 ± 0.39] x 10^−4^; p = 0.7; t-test). The strength of the responses of contralateral and ipsilateral eyes also remained the same in the dominant CPP treated animals after 4d MD. However, the response of the ipsilateral eye in submissive CPP-treated mice was significant increased after the 4d of MD ([1.98 ± 0.16] x 10^−4^ to [2.34 ± 0.17] x 10^−4^; p ≤ 0.05; t-test). But as the response of the contralateral eye also had a tendency to increase, the ODI of the submissive CPP-treated mice was not affected.

### 3.5 Social rank is independent from serotonin, GABA and LTP

During the administration of drugs that prevented OD plasticity, running wheel use was monitored on a daily basis (Fig. 4). Although the drugs had some global effects on behaviour and, consequently, motivation to enter the running wheel, neither WAY-100635 (Fig. 4A) nor diazepam (Fig. 4B) nor CPP (Fig 4C) changed the social hierarchy between the two mice of a pair.

**Figure 4:**
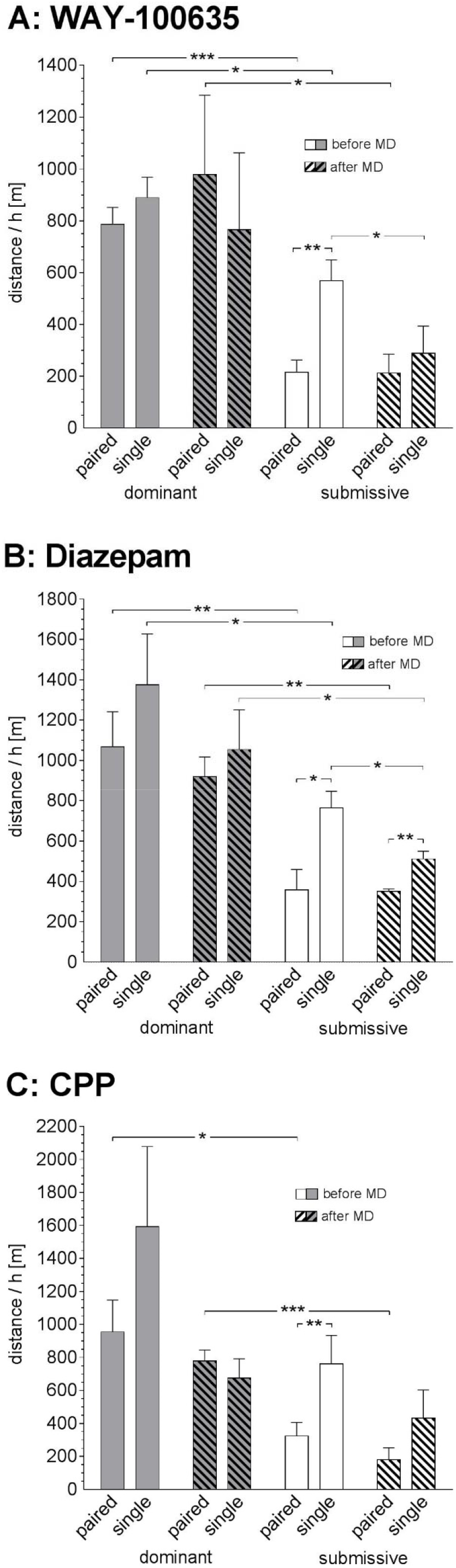
Running wheel data under WAY-100635-, diazepam- and CPP-treatment. Neither WAY-100635 (A) nor diazepam (B) nor CPP (C) changed the social hierarchy between the two mice of a pair.

As a serotonin antagonist, WAY-100635 predictably increased the aggression especially of the dominant mouse. In consequence, the dominant animal defended its access to the running wheel, whereas the submissive partner appeared even more subdued after the initiation of the WAY-100635 treatment. ANOVA confirmed an effect of dominance (F_1.12_ = 15.937, p ≤ 0.01).

In contrast, both diazepam and CPP decreased running wheel use of dominant and submissive mice alike (repeated measures factor, diazepam: F_1.12_ = 3.432, p = 0.089; CPP: F_1.14_ = 5.537, p ≤ 0.05, ANOVA), but social dominance remained the most influential factor (diazepam: F_1.12_ = 21.948, p ≤ 0.001; CPP: F_1.14_ = 12.656, p ≤ 0.01). Thus, when together, dominant mice kept running more than submissive ones in all treatment groups, with the latter turning the wheel more eagerly when alone.

These observations indicate that interference with serotonin, GABA or glutamate-NMDA transmission do not act on the social status to abolish OD plasticity in dominant mice. Rather, they work independent from the neuronal representation of social hierarchy, presumably locally in the visual cortex.

### 3.6 Dopamine regulates social dominance and ocular dominance plasticity of adult male mice

As the medial prefrontal cortex (mPFC) bidirectionally regulates the social status of mice (Wang et al. 2011), and can also exert an influence on V1 processing (Nguyen et al. 2015; Noudoost and Moore 2011), we wondered whether it might have the capacity to control ODP as well. Therefore, we first checked whether the visual cortex received input from the mPFC by performing retrograde tracing from anterior and posterior portions of the binocular visual cortex (Fig. 5A). Indeed, labeled neurons were found within deep layers in the anterior cingulate subregion of the mPFC. It is interesting to note that these projections appear to be topographically arranged, with more ventrally situated neurons sending axons to more caudal subregions of the visual cortex.

**Figure 5:**
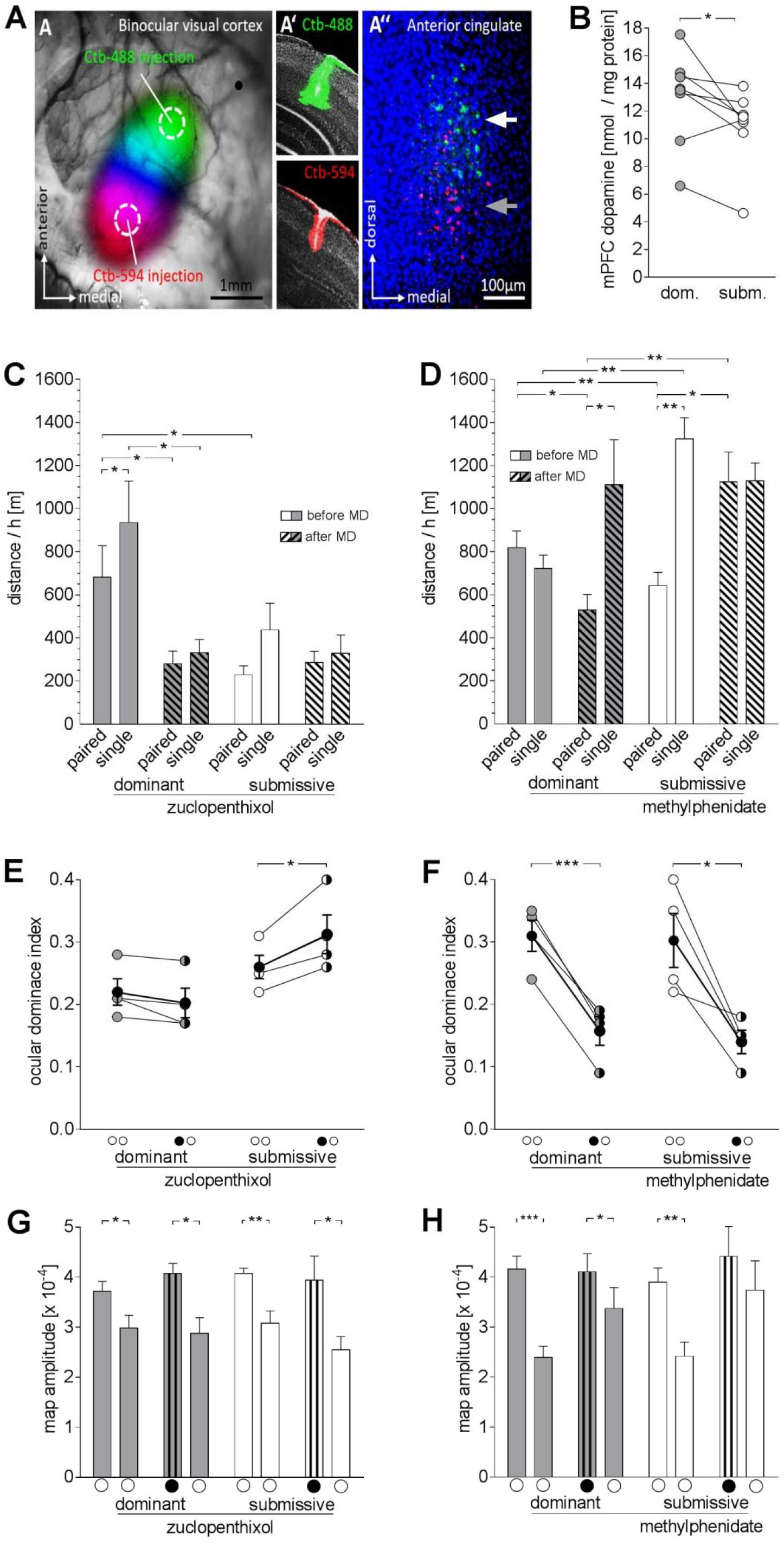
Cortical dopamine transmission regulates OD plasticity in mice. (A) By retrograde tracing of the visual cortex, labeled neurons were found in the mPFC. Anterior parts of V1 (green, brighter in black & white reproductions) are innervated by more dorsal parts of the anterior cingulate (white arrow), more posterior parts (red, darker in b/w) by the more ventral anterior cingulate (grey arrow). (B) Dopamine content is higher in the mPFC of dominant than submissive animals. (C) Zuclopenthixol treatment abolished the differential running wheel use of dominant and submissive mice. (D) The dominance relationship between the mice is partly reversed upon methylphenidate treatment. (E) The dopamine receptor antagonist zuclopenthixol blocked OD plasticity of the dominant mice and, as expected, of submissive mice as well. (F) In contrast, the dopaminergic agonist methylphenidate, administered to the submissive animal, resulted in both animals displaying OD plasticity. (G, H) Cortical response amplitudes elicited by stimulation of the contralateral and ipsilateral eyes. All conventions are as in Fig. 1.

Next, we determined the dopamine content in the mPFC by HPLC (Fig. 5B). In seven out of eight cases, the content was higher in the dominant than the submissive animal of a pair. On average, dominant mice had 12.96 ± 1.18 nmol/mg protein, and submissive mice had 10.93 ± 0.97 nmol/mg protein dopamine in the mPFC, a difference that proved to be significant (p ≤ 0.05, paired t-test). Thus, mPFC dopamine might play a role in mediating between social dominance and ocular dominance plasticity.

To check this assumption, we bidirectionally manipulated cortical dopamine transmission. In order to reduce it, we administered a low dose (0.2 mg/kg) of the neuroleptic zuclopenthixol. Zuclopenthixol blocks both D1 and D2 receptors, with little cross-action at other aminergic receptors. Moreover, the small dose is reported to reduce aggression, but to have no effect on motor behaviour, thus probably acting in the mPFC rather than the striatum (Manzaneque and Navarro 1999).

Indeed, this intervention not only reduced locomotion globally (F_1.6_ = 10.398, p ≤ 0.05, ANOVA) – as would be expected from a neuroleptic –, but specifically abolished the differential running wheel use of dominant and submissive mice (Fig. 5C). According to two-factor ANOVA with repeated measures, there was no overall effect of social dominance (F_1.6_ = 3.993, p > 0.05), but an interaction of social dominance with treatment (F_1.6_ = 8.439, p ≤ 0.05), which could be traced to significantly reduced running in dominant mice only after zuclopenthixol treatment, using post-hoc testing (p ≤ 0.05 for both paired and single running, paired t-tests). Running activity in submissive mice, in contrast, was in no way influenced by the neuroleptic.

Accordingly, optical imaging of intrinsic signals showed that zuclopenthixol treatment blocked OD plasticity of dominant animals (Fig. 5E, 0.22 ± 0.02 vs. 0.20 ± 0.02, n = 4), and even resulted in increased ODIs in submissive mice after 4d MD (0.26 ± 0.02 to 0.31 ± 0.03, n = 4). OD was not affected by either social rank (F_1.6_ = 5.118, p > 0.05) or MD (F_1.6_ = 5.345, p > 0.05), according to ANOVA, but there was an interaction (F_1.6_ = 21.382, p ≤ 0.01), which was due to the significantly increased ODI in submissive animals (p ≤ 0.05, paired t-test). The amplitude data (Fig. 5G) could offer no conclusive explanation for this unexpected change, since, apart from the striking influence of eye, no other factor had a significant influence, according to ANOVA.

Finally, we systemically administrated the dopamine reuptake inhibitor methylphenidate hydrochloride to submissive paired arena mice. Here, we faced the problem that social status is a relative concept, such that there cannot logically be two (out of two) dominant mice. Indeed, preliminary experiments had shown that administration of the drug to mice confined to the narrow cage, as well as administration to both mice in the arena, resulted in escalating violence that could not be tolerated. We therefore resorted to the arena with running-wheel paradigm established above, where it was possible for both mice to keep a distance. Here, we injected only the submissive mouse with a low dose of methylphenidate that has been shown to increase mPFC, but not striatal dopamine (Koda et al. 2010).

The dominance relationship between the mice indeed became more ambiguous, appearing partly reversed, upon this treatment (Fig. 5D). The methylphenidate-treated submissive mice increased their wheel-use at the expense of the dominant partners, who, in turn, resorted to more vigorous running when alone. While this pattern suggests a complete reversal of social hierarchy, behavioural observation showed a lot of agonistic behaviour and competition for the wheel that is more in line with the notion that the rank question remained unsettled during the treatment.

Remarkably, methylphenidate treatment of the submissive mouse indeed resulted in OD plasticity in both the originally dominant and the submissive mouse (Fig. 5F). As expected, the untreated dominant mice showed a strong shift of the ocular dominance towards the open, ipsilateral eye (0.31 ± 0.03 to 0.16 ± 0.02, n = 4). Amazingly, the methylphenidate-treated submissive mice also did show an OD shift of similar magnitude (0.30 ± 0.04 to 0.14 ± 0.02, n = 4). Thus, whereas MD had a highly significant influence on the data (F_1.6_ = 39.758, p ≤ 0.001), social dominance had none (F_1.6_ = 0.115, p > 0.5), and the shifts were significant in both groups, according to post-hoc paired t-tests. As the shift did not result in a balanced ODI of 0, deprived eye responses (fig. 5H) remained higher than open-eye responses in all cases (dominant: contra: [4.16 ± 0.26] x 10^−4^ vs. [4.11 ± 0.36] x 10^−4^, p = 0.9, ipsi: [2.39 ± 0.22] x 10^−4^ vs. [3.37 ± 0.12] x 10^−4^, p = 0.1, t-test; submissive: contra: [3.9 ± 0.28] x 10^−4^ vs. [4.42 ± 0.68] x 10^−4^, p = 0.4; ipsi: [2.42 ± 0.28] x 10^−4^ vs. [3.74 ± 0.58] x 10^−4^; p = 0.09; t-test), although there was an interaction of MD x eye (F_1.6_ = 48.148, p ≤ 0.001, ANOVA).

## 4. Discussion

In this paper we have shown the connection between social dominance status and ocular dominance plasticity: Whereas ocular dominance plasticity (ODP) can be observed in either mouse if two animals are housed together in a large arena (Balog et al. 2014), constraining the space or introducing a limited resource (running wheel) abolishes ODP in the submissive animal.

### Methodological considerations

We have based our assessment of relative social status mostly on the differential use of the running wheel. Observations of agonistic behaviour and physical appearance corroborated our judgement by this new measure, which was introduced for pragmatic reasons: Adult male mice that tolerate each other for a sufficient period of time to induce ODP (i.e., at least 4d) show very little agonistic behaviour to rely on (Balog et al. 2014). Even preliminary attempts to use a variant of the tube test (Malatynska et al. 2002) failed to yield conclusive results. A running wheel, however, has been shown to be a highly attractive feature for mice (Meijer and Robbers 2014). By comparing its use when mice were together and when alone, we could make sure that lesser use in one animal was not due to lack of interest. In addition, this procedure provided the submissive mouse with some running time. Although we have previously conclusively shown that wheel running over only a few days of monocular deprivation (MD) is ineffective in inducing ODP in adult mice (Balog et al. 2014), this arrangement makes such a causation additionally unlikely in the present study.

Nonetheless, another study (Kalogeraki et al. 2014) has claimed to have induced OD plasticity in fully adult male mice by providing access to a running wheel during seven days of MD. In that study, however, the mice where apparently housed socially (three to five mice per cage). We have shown that social housing induced OD plasticity (Balog et al. 2014), but a control group of socially housed animals without running wheel is lacking in the conflicting study. Moreover, they used seven days of MD, whereas we used four days, which is the standard duration to check for critical-period like plasticity (Gordon and Stryker 1996). In our previous study, we showed that providing single housed mice with a running wheel throughout four days of MD failed to induce OD plasticity (Balog et al. 2014). We here substantiate this observation by finding no correlation between running distance and plasticity in neither dominant nor submissive mice. Finally, as stated in the introduction, the differing OD plasticity among two male mice housed together in a standard cage was already observed in our first study – sparking the investigation presented here -, although no running wheel was present. Thus, we can safely exclude that our method to assess social dominance could in any way interfere with OD plasticity.

### OD plasticity was Hebbian and required attenuated GABA and increased serotonin transmission

ODP in dominant mice was shown to be Hebbian in the present study, as it was blocked by the NMDA antagonist CPP, thus confirming the current knowledge on the early phase of ODP (Sawtell et al. 2003; Sato and Stryker 2008; Ranson et al. 2012). Furthermore, it required reduced GABA inhibition and increased serotonin transmission. This, too, is in line with a host of studies (Sale et al. 2007; Maya-Vetencourt et al. 2008; 2011; Spolidoro et al. 2009; Harauzov et al. 2011). In addition, serotonin is strongly involved in the regulation of social behaviour, and its release is altered by housing conditions (Miura et al. 2008). However, HPLC of the visual cortices of the animals we examined did not indicate that the concentrations and ratio (turnover) of serotonin and its metabolite 5-HIAA differed between dominant and submissive mice. However, serotonin turnover in paired mice – both with and without a running wheel – was lower than in non-plastic single house mice, and ocular dominance plasticity of dominant mice could be blocked by diazepam and WAY-100635. Thus, it may be assumed that changes in GABA and serotonin activity, although not sufficient, are necessary to allow adult OD plasticity. It follows that they do not act downstream of the mechanisms that differentially regulate OD plasticity in dominant and submissive mice. Seeing that in arena mice, introduction of a running wheel – a very popular and competitive resource – blocks plasticity in submissive mice where it would otherwise be present, we suggest that social housing of mice enables OD plasticity by mechanisms that involve the serotonin and GABA systems (Balog et al. 2014). This plasticity can then, however, secondarily be blocked along an independent route by the experience of submissiveness. In line with this reasoning, the treatments of diazepam, WAY-100635 and CPP did not affect the dominant and submissive behaviour of the mice. Thus, ocular dominance plasticity mediated by social status has a different origin than the primary visual cortex.

### Higher-order cortices are involved in regulating adult OD plasticity

In line with this reasoning, we have shown that manipulation of dopamine transmission, using dopaminergic drugs at concentrations that have no effects in the striatum, but in the PFC (Manzaneque and Navarro 1999; Koda et al. 2010), bidirectionally regulated ODP. As dopamine afferents are only received by higher order cortices, but spare the primary sensory cortices completely, this observation proves that plasticity in a primary sensory cortex is controlled by higher-order cortices, most likely the medial PFC (mPFC), which is a key structure for cognition, mental health, and social behaviour (Wang et al. 2011, 2014; Wass et al. 2018). Social status is bidirectionally regulated by the mPFC (Wang et al. 2011). In rodents, the PFC receives an exclusive projection of dopamine fibres from the ventral tegmentum, which are prominently involved in its functioning (Winterfeld et al. 1998; Wass et al. 2018), including the establishment of social hierarchy (Yamaguchi et al. 2017). By its efferents, the mPFC can exert widespread influences on brain functions. I.a., it has direct connections to the visual cortex (Sesack et al. 1989; Zhang et al. 2016), which we have confirmed in the present study and shown to be topographic in nature. By these connections, the mPFC can regulate sensory processing and attention in the visual cortex (Zhang et al. 2014; Nguyen et al. 2015; Noudoost and Moore, 2011).

In accordance with these studies, we have shown in the present study that dopamine content in the mPFC was higher in dominant than submissive animals. Using both a specific agonist and antagonist, we could further demonstrate that the dominance behaviour of the mice is bidirectionally altered by dopamine transmission, resulting in corresponding changes in the ocular dominance plasticity within the primary visual cortex. The fact that, unlike serotonergic, GABAergic or glutamatergic interventions described above, dopaminergic interventions interfered not only with ODP, but also with the behavioural expression of social status, strongly argues for a causal link between these phenomena. I.e., whereas certain levels of serotonergic and GABAergic transmission are merely permissive for adult ODP, a certain level of dopaminergic transmission in the mPFC is necessary.

It remains an open question how the mPFC exerts its influence on V1 plasticity. Although there is a prominent projection from the mPFC to the dorsal raphe nuclei (Sesack et al 1989; Peyron et al. 1998; Vázquez-Borsetti et al. 2009), we can exclude this route, as serotonin content and turnover were similar in dominant and submissive mice. We rather favour a direct cortico-cortical connection, the existence of which is established (Sesack et al. 1989; Zhang et al. 2016), has been confirmed in the present study, and is known to regulate visual attention (Noudost and Moore 2011). This suggestion can be further substantiated by other studies that have already shown that other cortical areas also affect processing and plasticity in the primary visual cortex (Teichert and Bolz, 2017; Teichert et al. 2018a, 2018b, 2018c).

### Additional observations

Two unexpected observations during the course of this study merit some brief remarks. First, WAY-100635 treatment not only blocked plasticity, but resulted in even increased ODIs after MD. This ODI increase is due to diminished response strength of the ipsilateral eye in the WAY 100635-treated mice after 4d of MD. In a complementary fashion, we recently observed an augmentation of ipsilateral response strength upon auditory or whisker deprivation (Teichert et al. 2019) that was accompanied by an increased serotonin turnover in the visual cortex and could be blocked by WAY 100635 (additional unpublished observations). Taken together, these results tentatively suggest that higher serotonin transmission increases and less of it decreases the cortical response to the ipsilateral eye, although it remains mysterious how this is brought about.

Second, it was also surprising that, while the ODI shift in dominant animals was uniform in all control groups, the underlying mechanism differed markedly: In untreated animals and animals with a single daily vehicle injection, it was achieved by an increase of ipsilateral response (so-called adult plasticity, Ranson et al. 2012; Sato and Stryker 2008). If mice received two daily vehicle injections, however, a decrease of the contralateral eye response was observed – so called juvenile plasticity. This indicates that the brain can switch rather easily between these two modes of plasticity, both of which are basically Hebbian and appear to be distinguished only by the sliding threshold between long-term depression and potentiation (Philpot et al. 2003). In the present case, the switch might have been effected by the different stress levels caused by the injection regime. In line with these considerations, small differences in the housing procedure might also explain why we here observed adult plasticity in untreated arena mice (unrelated, group housed males) without running wheel, but documented juvenile plasticity in our previous study (previously isolated brothers, Balog et al. 2014).

### Conclusion

We have shown in this study that ODP in adult male mice is dependent on social status. We have made a strong case for the assumption that the mPFC, which represents the social status, is on the one hand influenced by dopamine transmission to organize submissive or dominant patterns of behaviour, and on the other hand suppresses ODP in the visual cortex of submissive animals, probably via a direct projection. The idea that associative cortices might regulate plasticity in the primary cortices has been uttered before (He et al. 2006, 2007; Lehmann 2010), but has, to our knowledge, never as yet been experimentally tested. Future studies using local application of drugs and optogenetic manipulations could show how this influence is mediated on a functional level.

## 5. Acknowledgements

We are obliged to Prof. Jürgen Bolz for constant support and helpful discussion. Thanks are further due to Elisabeth Meier for excellent technical assistance and Sandra Eisenberg for animal care. Finally, we wish to thank Dr. John O’Ball for proof-reading the manuscript. Jenny Balog was supported by a Landesgraduiertenstipendium during the preparation of this study.

## References

Balog J, Matthies U, Naumann L, Voget M, Winter C, Lehmann K (2014) Social experience modulates ocular dominance plasticity differentially in adult male and female mice. Neuroimage. 103:454–461. doi: 10.1016/j.neuroimage.2014.08.040.

Baroncelli L, Sale A, Viegi A, Maya Vetencourt JF, De Pasquale R, Baldini S, Maffei L (2010) Experience-dependent reactivation of ocular dominance plasticity in the adult visual cortex. Exp. Neurol. 226, 100–109.

Blanchard DC, Spencer RL, Weiss SM, Blanchard RJ, McEwen B, Sakai RR (1995) Visible burrow system as a model of chronic social stress: behavioral and neuroendocrine correlates. Psychoneuroendocrinology. 20:117–34.

Cang J, Kalatsky VA, Löwel S, Stryker MP, (2005) Optical imaging of the intrinsic signal as a measure of cortical plasticity in the mouse. Vis Neurosci. 22:685–91.

Colas-Zelin D, Light KR, Kolata S, Wass C, Denman-Brice A, Rios C, Szalk K, Matzel LD (2012) The imposition of, but not the propensity for, social subordination impairs exploratory behaviors and general cognitive abilities. Behav Brain Res 232(1): 294–305.

Desjardins JK, Fernald RD (2008) How do social dominance and social information influence reproduction and the brain? Integr Comp Biol. 48:596–603. doi: 10.1093/icb/icn089.

Enard W, Gehre S, Hammerschmidt K, Hölter SM, Blass T, Somel M, Brückner MK, Schreiweis C, Winter C, Sohr R, Becker L, Wiebe V, Nickel B, Giger T, Müller U, Groszer M, Adler T, Aguilar A, Bolle I, Calzada-Wack J, Dalke C, Ehrhardt N, Favor J, Fuchs H, Gailus-Durner V, Hans W, Hölzlwimmer G, Javaheri A, Kalaydjiev S, Kallnik M, Kling E, Kunder S, Mossbrugger I, Naton B, Racz I, Rathkolb B, Rozman J, Schrewe A, Busch DH, Graw J, Ivandic B, Klingenspor M, Klopstock T, Ollert M, Quintanilla-Martinez L, Schulz H, Wolf E, Wurst W, Zimmer A, Fisher SE, Morgenstern R, Arendt T, de Angelis MH, Fischer J, Schwarz J, Pääbo S (2009) A humanized version of Foxp2 affects cortico-basal ganglia circuits in mice. Cell. 137:961–71. doi: 10.1016/j.cell.2009.03.041.

Felice LJ, Felice JD, Kissinger PT (1978) Determination of catecholamines in rat brain parts by reverse-phase ion-pair liquid chromatography. J Neurochem. 31:1461–5.

Fitchett AE, Collins SA, Barnard CJ, Cassaday HJ (2005) Subordinate male mice show long-lasting differences in spatial learning that persist when housed alone. Neurobiol Learn Mem. 84:247–51.

Fletcher A, Forster EA, Bill DJ, Brown G, Cliffe IA, Hartley JE, Jones DE, McLenachan A, Stanhope KJ, Critchley DJ, Childs KJ, Middlefell VC, Lanfumey L, Corradetti R, Laporte AM, Gozlan H, Hamon M, Dourish CT (1996) Electrophysiological, biochemical, neurohormonal and behavioural studies with WAY-100635, a potent, selective and silent 5-HT1A receptor antagonist. Behav Brain Res. 73:337–53.

Forster EA, Cliffe IA, Bill DJ, Dover GM, Jones D, Reilly Y, Fletcher A (1995) A pharmacological profile of the selective silent 5-HT1A receptor antagonist, WAY-100635. Eur J Pharmacol. 281:81–8.

Giovanoli S, Engler H, Engler A, Richetto J, Voget M, Willi R, Winter C, Riva MA, Mortensen PB, Feldon J, Schedlowski M, Meyer U (2013) Stress in puberty unmasks latent neuropathological consequences of prenatal immune activation in mice. Science. 339:1095–9. doi: 10.1126/science.1228261.

Goeckner DJ, Greenough WT, Mead WR (1973) Deficits in learning tasks following chronic overcrowding in rats. J Pers Soc Psychol. 28:256–61.

Gordon JA, Stryker MP (1996) Experience-dependent plasticity of binocular responses in the primary visual cortex of the mouse. J Neurosci. 16:3274–86.

Hanover JL, Huang ZJ, Tonegawa S, Stryker MP (1999) Brain-derived neurotrophic factor overexpression induces precocious critical period in mouse visual cortex. J Neurosci. 19:RC40.

Harauzov A, Spolidoro M, DiCristo G, De Pasquale R, Cancedda L, Pizzorusso T, Viegi A, Berardi N, Maffei L (2010) Reducing intracortical inhibition in the adult visual cortex promotes ocular dominance plasticity. J Neurosci. 30:361–71. doi: 10.1523/JNEUROSCI.2233-09.2010.

He HY, Hodos W, Quinlan EM (2006) Visual deprivation reactivates rapid ocular dominance plasticity in adult visual cortex. J Neurosci. 26:2951–5.

He HY, Ray B, Dennis K, Quinlan EM (2007) Experience-dependent recovery of vision following chronic deprivation amblyopia. Nat Neurosci. 10:1134–6.

Huang ZJ, Kirkwood A, Pizzorusso T, Porciatti V, Morales B, Bear MF, Maffei L, Tonegawa S (1999) BDNF regulates the maturation of inhibition and the critical period of plasticity in mouse visual cortex. Cell. 98:739–55.

Jetz W, Rubenstein DR (2011) Environmental uncertainty and the global biogeography of cooperative breeding in birds. Curr Biol. 21:72–8. doi: 10.1016/j.cub.2010.11.075.

Kalatsky VA, Stryker MP (2003). New paradigm for optical imaging: temporally encoded maps of intrinsic signal. Neuron. 38(4):529–45.

Kalogeraki E, Greifzu F, Haack F, Löwel S (2014) Voluntary physical exercise promotes ocular dominance plasticity in adult mouse primary visual cortex. J. Neurosci. 34(46): 15476–15481.

Kappel S, Hawkins P, Mendl MT (2017) To Group or Not to Group? Good Practice for Housing Male Laboratory Mice. Animals (Basel). 7. pii: E88. doi: 10.3390/ani7120088

Kar F, Whiting MJ, Noble DWA (2017) Dominance and social information use in a lizard. Anim Cogn. 20:805–812. doi: 10.1007/s10071-017-1101-y.

Koda K, Ago Y, Cong Y, Kita Y, Takuma K, Matsuda T (2010) Effects of acute and chronic administration of atomoxetine and methylphenidate on extracellular levels of noradrenaline, dopamine and serotonin in the prefrontal cortex and striatum of mice. J Neurochem. 114:259–70. doi: 10.1111/j.1471-4159.2010.06750.x.

Lehmann K, Löwel S (2008) Age-dependent ocular dominance plasticity in adult mice. PLoS One 3, e3120.

Lehmann K (2010) Gemeinsamkeiten und Unterschiede in der Entwicklungsplastizität von assoziativen und primärsensorischen Kortexgebieten. Habilitationsschrift, Friedrich Schiller-Universität Jena.

Lehmann K, Schmidt KF, Löwel S (2012) Vision and visual plasticity in ageing mice. Restor Neurol Neurosci. 30:161–78. doi: 10.3233/RNN-2012-110192.

Malatynska E, Goldenberg R, Shuck L, Haque A, Zamecki P, Crites G, Schindler N, Knapp RJ (2002) Reduction of submissive behavior in rats: a test for antidepressant drug activity. Pharmacology 64: 8–17.

Manzaneque JM, Navarro JF (1999) An ethopharmacological assessment of the effects of zuclopenthixol on agonistic interactions in male mice. Methods Find Exp Clin Pharmacol. 21:11–5.

Matzel LD, Kolata S, Light K, Sauce B (2017) The tendency for social submission predicts superior cognitive performance in previously isolated male mice. Behav Processes. 134:12–21. doi: 10.1016/j.beproc.2016.07.011.

Maya Vetencourt JF, Sale A, Viegi A, Baroncelli L, De Pasquale R, O’Leary OF, Castrén E, Maffei L (2008) The antidepressant fluoxetine restores plasticity in the adult visual cortex. Science. 320: 385–388.

Maya Vetencourt JF, Tiraboschi E, Spolidoro M, Castrén E, Maffei L (2011) Serotonin triggers a transient epigenetic mechanism that reinstates adult visual cortex plasticity in rats. Eur J Neurosci. 33: 49–57. doi: 10.1111/j.1460-9568.2010.07488.x.

Meijer JH, Robbers Y (2014) Wheel running in the wild. Proc Biol Sci. 281. pii: 20140210. doi: 10.1098/rspb.2014.0210.

Morgan D, Grant KA, Gage HD, Mach RH, Kaplan JR, Prioleau O, Nader SH, Buchheimer N, Ehrenkaufer RL, Nader MA (2002) Social dominance in monkeys: dopamine D2 receptors and cocaine self-administration. Nat Neurosci. 5:169–74.

Nguyen HN, Huppé-Gourgues F, Vaucher E (2015) Activation of the mouse primary visual cortex by medial prefrontal subregion stimulation is not mediated by cholinergic basalo-cortical projections. Front Syst Neurosci. 9:1. doi: 10.3389/fnsys.2015.00001.eCollection

Miura H, Ozaki N, Shirokawa T, Isobe K (2008) Changes in brain tryptophan metabolism elicited by ageing, social environment, and psychological stress in mice. Stress. 11:160–9. doi: 10.1080/10253890701685908.

Noudoost B, Moore T (2011) Control of visual cortical signals by prefrontal dopamine. Nature. 474:372–5. doi: 10.1038/nature09995.

Olsson IA, Sherwin CM (2006) Behaviour of laboratory mice in different housing conditions when allowed to self-administer an anxiolytic. Lab Anim. 40:392–9.

Paxinox G, Franklin K (2012) The Mouse Brain in Stereotaxic Coordinates. Academic Press, Australia

Peyron C, Petit JM, Rampon C, Jouvet M, Luppi PH (1998) Forebrain afferents to the rat dorsal raphe nucleus demonstrated by retrograde and anterograde tracing methods. Neuroscience. 82:443–68.

Philpot BD, Espinosa JS, Bear MF (2003) Evidence for altered NMDA receptor function as a basis for metaplasticity in visual cortex. J Neurosci. 23:5583–8.

Prabhu VV, Nguyen TB, Cui Y, Oh YE, Lee KH, Bagalkot TR, Chung YC (2018) Effects of social defeat stress on dopamine D2 receptor isoforms and proteins involved in intracellular trafficking. Behav Brain Funct. 14:16. doi: 10.1186/s12993-018-0148-5.

Ranson A, Cheetham CE, Fox K, Sengpiel F (2012) Homeostatic plasticity mechanisms are required for juvenile, but not adult, ocular dominance plasticity. Proc Natl Acad Sci USA. 109:1311–6. doi: 10.1073/pnas.1112204109.

Sale A, Maya Vetencourt JF, Medini P, Cenni MC, Baroncelli L, De Pasquale R, Maffei L (2007) Environmental enrichment in adulthood promotes amblyopia recovery through a reduction of intracortical inhibition. Nat Neurosci. 10:679–81. Epub 2007 Apr 29.

Sarna JR, Dyck RH, Whishaw IQ (2000) The Dalila effect: C57BL6 mice barber whiskers by plucking. Behav Brain Res. 108:39–45.

Sato M, Stryker MP (2008) Distinctive features of adult ocular dominance plasticity. J Neurosci. 28:10278–86. doi: 10.1523/JNEUROSCI.2451-08.2008.

Sawtell NB, Frenkel MY, Philpot BD, Nakazawa K, Tonegawa S, Bear MF (2003) NMDA receptor-dependent ocular dominance plasticity in adult visual cortex. Neuron. 38:977–85.

Sbragaglia V, Leiva D, Arias A, Antonio García J, Aguzzi J, Breithaupt T (2017) Fighting over burrows: the emergence of dominance hierarchies in the Norway lobster (Nephrops norvegicus). J Exp Biol. 220:4624–4633. doi: 10.1242/jeb. 165969.

Sesack SR, Deutch AY, Roth RH, Bunney BS (1989) Topographical organization of the efferent projections of the medial prefrontal cortex in the rat: an anterograde tract-tracing study with Phaseolus vulgaris leucoagglutinin. J Comp Neurol. 290:213–42.

Sperk G, Berger M, Hörtnagl H, Hornykiewicz O (1981) Kainic acid-induced changes of serotonin and dopamine metabolism in the striatum and substantia nigra of the rat. Eur J Pharmacol. 74:279–86.

Sperk G (1982) Simultaneous determination of serotonin, 5-hydroxindoleacetic acid, 3,4-dihydroxyphenylacetic acid and homovanillic acid by high performance liquid chromatography with electrochemical detection. J Neurochem. 38:840–3.

Spolidoro M, Sale A, Berardi N, Maffei L (2009) Plasticity in the adult brain: lessons from the visual system. Exp Brain Res. 192:335–41. doi: 10.1007/s00221-008-1509-3.

Spritzer MD, Meikle DB, Solomon NG (2004) The relationship between dominance rank and spatial ability among male meadow voles (Microtus pennsylvanicus). J Comp Psychol. 118(3): 332–9.

Stears K, Kerley GI, Shrader AM (2014) Group-living herbivores weigh up food availability and dominance status when making patch-joining decisions. PLoS One. 9:e109011. doi: 10.1371/journal.pone.0109011.

Stodieck SK, Greifzu F, Goetze B, Schmidt KF, Löwel S (2014) Brief dark exposure restored ocular dominance plasticity in aging mice and after a cortical stroke. Exp Gerontol. 60:1–11. doi: 10.1016/j.exger.2014.09.007.

Teichert M, Bolz J (2017) Simultaneous intrinsic signal imaging of auditory and visual cortex reveals profound effects of acute hearing loss on visual processing. Neuroimage. 159:459–472. doi: 10.1016/j.neuroimage.2017.07.037.

Teichert M, Isstas M, Wenig S, Setz C, Lehmann K, Bolz J (2018a) Cross-modal refinement of visual performance after brief somatosensory deprivation in adult mice. Eur J Neurosci. 47:184–191. doi: 10.1111/ejn.13798.

Teichert M, Isstas M, Zhang Y, Bolz J (2018b) Cross-modal restoration of ocular dominance plasticity in adult mice. Eur J Neurosci. 47:1375–1384. doi: 10.1111/ejn.13944.

Teichert M, Isstas M, Wieske F, Winter C, Bolz J (2018c) Cross-modal Restoration of Juvenile-like Ocular Dominance Plasticity after Increasing GABAergic Inhibition. Neuroscience. 393:1–11. doi: 10.1016/j.neuroscience.2018.09.040.

Teichert M, Isstas M, Liebmann L, Hübner CA, Wieske F, Winter C, Lehmann K, Bolz J (2019) Visual deprivation independent shift of ocular dominance induced by cross-modal plasticity. PLoS One 14(3): e0213616.

Vázquez-Borsetti P, Cortés R, Artigas F (2009) Pyramidal neurons in rat prefrontal cortex projecting to ventral tegmental area and dorsal raphe nucleus express 5-HT2A receptors. Cereb Cortex. 19:1678–86. doi: 10.1093/cercor/bhn204

Villarreal DM, Do V, Haddad E, Derrick BE (2002) NMDA receptor antagonists sustain LTP and spatial memory: active processes mediate LTP decay. Nat Neurosci. 5:48–52.

Wang F, Zhu J, Zhu H, Zhang Q, Lin Z, Hu H (2011) Bidirectional control of social hierarchy by synaptic efficacy in medial prefrontal cortex. Science. 334:693–7. doi: 10.1126/science.1209951.

Wang F, Kessels HW, Hu H (2014) The mouse that roared: neural mechanisms of social hierarchy. Trends Neurosci. 37:674–82. doi: 10.1016/j.tins.2014.07.005.

Wass C, Sauce B, Pizzo A, Matzel LD (2018) Dopamine D1 receptor density in the mPFC responds to cognitive demands and receptor turnover contributes to general cognitive ability in mice. Sci Rep. 8:4533. doi: 10.1038/s41598-018-22668-0.

Winter C, Djodari-Irani A, Sohr R, Morgenstern R, Feldon J, Juckel G, Meyer U (2009) Prenatal immune activation leads to multiple changes in basal neurotransmitter levels in the adult brain: implications for brain disorders of neurodevelopmental origin such as schizophrenia. Int. J. Neuropsychopharmacol. 12, 513–524.

Winterfeld KT, Teuchert-Noodt G, Dawirs RR (1998) Social environment alters both ontogeny of dopamine innervation of the medial prefrontal cortex and maturation of working memory in gerbils (Meriones unguiculatus). J Neurosci Res. 52:201–9.

Yamaguchi Y, Lee YA, Kato A, Goto Y (2017) The Roles of Dopamine D1 Receptor on the Social Hierarchy of Rodents and Nonhuman Primates. Int J Neuropsychopharmacol. 20(4):324–335. doi: 10.1093/ijnp/pyw106.

Zhang S, Xu M, Kamigaki T, Hoang Do JP, Chang WC, Jenvay S, Miyamichi K, Luo L, Dan Y (2014) Selective attention. Long-range and local circuits for top-down modulation of visual cortex processing. Science. 345:660–5. doi: 10.1126/science.1254126.

Zhang S, Xu M, Chang WC, Ma C, Hoang Do JP1, Jeong D, Lei T, Fan JL, Dan Y (2016) Organization of long-range inputs and outputs of frontal cortex for top-down control. Nat Neurosci. 19:1733–1742. doi: 10.1038/nn.4417.

